# Sex and estrous cycle stage shape left-right asymmetry in chronic hippocampal seizures in mice

**DOI:** 10.1101/2023.01.20.524965

**Authors:** Cathryn A. Cutia, Leanna K. Leverton, Catherine A. Christian-Hinman

## Abstract

Lateralization of hippocampal function is indicated by varied outcomes of patients with neurological disorders that selectively affect one hemisphere of this structure, such as temporal lobe epilepsy (TLE). The intrahippocampal kainic acid (IHKA) injection model of TLE allows for targeted damage to the left or right hippocampus, enabling systematic comparison of effects of left-right asymmetry on seizure and non-seizure outcomes. Although varying non-seizure phenotypic outcomes based on injection side in dorsal hippocampus were recently evaluated in this model, differences in chronic seizure patterns in left- (IHKA-L) vs. right-injected (IHKA-R) IHKA animals have yet to be evaluated. Here, we evaluated hippocampal seizure incidence in male and female IHKA-L and IHKA-R mice. Females displayed increased electrographic seizure activity compared to males at both 2 months and 4 months post-injection (mpi). In addition, IHKA-L females showed higher seizure frequency than IHKA-R on diestrus and estrus at 2 mpi, but seizure duration and time in seizures were only higher in IHKA-L females on diestrus. These cycle stage-associated changes, however, did not persist to 4 mpi. Furthermore, this lateralized difference in seizure burden was not observed in males. These results indicate for the first time that the side of IHKA injection can shape chronic electrographic seizure burden. Overall, these results demonstrate a female-specific left-right asymmetry in hippocampal function can interact with estrous cycle stage to shape chronic seizures in mice with epilepsy, with implications for neural activity and behavior in both normal and disease states.

## Introduction

The human brain is structurally and functionally lateralized.^1^ Although the hippocampus shows structural symmetry, neuroimaging studies suggest distinct functional roles of the two human hippocampi.^2,3^ Importantly, several neurological diseases affect the hippocampus. The damage inflicted from neurological disorders such as stroke and epilepsy can be unilaterally localized. Furthermore, patient outcomes can vary based on whether this damage is present in the left or right hippocampus. For example, patients with ischemic stroke damage in the left hippocampus have more apparent memory dysfunction than those with damage in the right.^4^ Additionally, people with epilepsy whose seizures arise from focal damage to the left hippocampus show a higher degree of cognitive impairment than with right hippocampal seizure foci.^5–7^ This variation in outcomes based on the hemisphere containing the unilateral damage may result from underlying functional lateralization in the hippocampus.

Temporal lobe epilepsy (TLE) is a common form of focal epilepsy in which seizures arise from a specific subregion and hemisphere of the temporal lobe, particularly the hippocampus.^8^ Interestingly, clinical observations in women with TLE suggest that the lateralization of a patient’s seizure focus can impact seizure patterning in relationship to the menstrual cycle. For instance, clinical evidence suggests left-sided seizure foci are associated with higher incidence of perimenstrual catamenial seizures, whereas right-sided seizure foci are associated with non-catamenial pattering of seizures spread across the menstrual cycle.^9,10^ Women with catamenial epilepsy are at higher risk for resistance to antiseizure medications,^11^ underscoring the importance of identifying the underlying mechanisms. However, little is known about the neural mechanisms linking seizure focus lateralization to differential seizure patterning.

Developments demonstrating lateralization in the rodent hippocampus^12^ support the use of rodent models in investigating the mechanisms that underlie these differences. The intrahippocampal kainic acid (IHKA) mouse model of TLE, which allows for epileptogenic insults to be selectively targeted to the left or right hemisphere, shows neuropathological changes similar to human TLE.^13–15^ This model also displays epileptiform discharges and spontaneous recurrent seizures^13^ that are hallmarks of human TLE.^16–18^ To date, researchers using this model have arbitrarily targeted the left or right hippocampus for KA injection, in the absence of systematic examination of differential outcomes of left and right IHKA injections. In a recent study, however, we compared non-seizure phenotypic outcomes in C57BL/6J females injected with KA in the left or the right dorsal hippocampus.^19^ We found that dentate gyrus granule cell dispersion was altered in an injection site-specific manner,^19^ indicating lateralized phenotypes at the level of the hippocampus. However, recent work has suggested that granule cell dispersion does not correlate to chronic seizure severity in left-injected IHKA mice.^20^ Therefore, whether the lateralization effect in granule cell dispersion contributes to functional differences in hippocampal seizure incidence remains unclear. Furthermore, it is unknown whether potential lateralized differences in seizure occurrence are shaped by animal sex and, in females, estrous cycle stage. Here, we tested the hypothesis that the laterality of IHKA injection leads to differential patterning of subsequent spontaneous recurrent seizures, both based on sex and estrous cycle stage.

## Methods

### Animals and estrous cycle monitoring

Animal procedures used in this study complied with the ARRIVE guidelines and were approved by the Institutional Animal Care and Use Committee of the University of Illinois Urbana-Champaign. Female and male C57BL/6J mice (#000664, Jackson Laboratories) were purchased for delivery at 6 weeks of age. Mice were then housed in a 14:10 h light:dark cycle (lights off at 1900 h) and given food and water *ad libitum*.

Beginning one week after arrival, estrous cycle monitoring in females was performed between 0900 and 1100 h using a vaginal cytology protocol previously described.^21^ Female mice were assessed for at least 14 days to verify regular cycles prior to entering the study and resumed estrous cycle monitoring for the duration of the LFP recording periods. To evaluate the development of differences in seizures over time, mice were recorded and underwent estrous cycle monitoring at both 2 and 4 mpi. Some female mice (IHKA-R = 2, IHKA-L = 5) included in this study were included in a previous publication,^22^ but the analysis of the data from these animals carried out in the previous work is distinct from that reported here. Additionally, the LFP data collected at 1 mpi from the female animals in the present study were included in another study, and are therefore not shown here.^19^

### Stereotaxic IHKA and LFP electrode implantation surgeries

All stereotaxic surgeries were carried out under isoflurane anesthesia (2-3%, vaporized in 100% oxygen) with carprofen (0.5 mg/ml) for analgesia. Female mice underwent stereotaxic unilateral injection of KA (Tocris Bioscience; 50 nl of 20 mM prepared in 0.9% sterile saline) on the first day of diestrus following the first estrous cycle monitoring period. Age-matched males (> postnatal day 60) were injected in the same manner. Injections were randomly targeted to the left or right dorsal hippocampal region (relative to Bregma: 2.0 mm posterior, 1.5 mm lateral, 1.4 mm ventral to cortical surface) as previously described.^23^ Age-matched controls were injected with the same volume at the same location with saline. Saline-injected animals (female: left n = 5, right n = 5; males: left n = 3, right n = 4) showed no seizures and thus were not included in analyses.

All animals were allowed two weeks to recover from the injection before undergoing a second surgery for LFP electrode implantation. Two twisted bipolar electrodes (P1 Technologies) were implanted into the ipsilateral hippocampus just dorsal and lateral to the injection site (relative to Bregma: 2.0 mm posterior, 1.75 mm lateral, 1.25 mm ventral).^19,22,24,25^ Anchor micro-screws (J.I. Morris Co.) were placed into the skull and stabilized with dental cement (Teets “Cold Cure” Dental Cement). Mice were singly housed after electrode implantation and for the duration of the remaining experimental period.

### LFP recording and analysis

One week after electrode implantation, the mice were tethered to an electrical commutator (P1 Technologies) connected to a Brownlee 440 amplifier (NeuroPhase) with gain set at 1 K. LFP signals were recorded as the local field potential differential between the two electrodes,^25^ sampled at 2 KHz and digitized to recorder software written in MATLAB.^25^ Mice were recorded at 1, 2, and 4 mpi. All recordings were analyzed with an automated electrographic seizure analyzer^26^ with the minimal seizure duration set at five seconds and the interictal interval at six seconds. The 2- and the 4-mpi recording periods were evaluated in the current study.

### Statistical analysis

All statistical comparisons were performed using R software. Comparisons between male and female IHKA-L and -R animals were made using two-way ANOVA and Tukey’s *post hoc* tests. Comparisons between IHKA-L and -R females within individual estrous cycle stages were made using two-sample t-tests, or Wilcoxon rank sum tests depending on the normality of the data. Normality was evaluated using Shapiro-Wilks tests, and homogeneity of variance was evaluated using Levene’s tests.

## Results

### Seizure burden is increased in IHKA females compared to males at 2 months after injection

Both male (IHKA-L = 9, IHKA-R = 8) and female (IHKA-L = 17, IHKA-R = 20) mice were evaluated to characterize sex differences in general seizure patterning and in response to varied side of targeted injection (**Figure 1A**). Recordings collected across a 7-day period at 2 mpi were evaluated for each mouse and averaged for group comparisons. Seizure frequency differed between groups based on both injection site (F(1,51) = 4.12, p = 0.047, **Figure 1B**) and sex (F(1,51) = 12.40, p = 0.0009, **Figure 1B**). Specifically, IHKA-L females displayed elevated seizure frequency compared to both IHKA-L (p = 0.005, Tukey, **Figure 1B**) and IHKA-R males (p = 0.002, **Figure 1B**). IHKA-R females also showed higher seizure frequency than IHKA-R males (p = 0.005). Seizure duration also displayed a difference based on sex (F(1,51) = 9.79, p = 0.002, **Figure 1C**), with IHKA-L females displaying longer seizure duration compared to both IHKA-L (p = 0.02, **Figure 1C**) and IHKA-R males (p = 0.03, **Figure 1C**). IHKA-R females also showed higher duration compared to IHKA-R males (p = 0.02, **Figure 1C**). The average percent time spent in seizures also differed by injection site (F(1,51) = 4.45, p = 0.04, **Figure 1D**) and sex (F(1,51) = 17.14, p = 0.0001, **Figure 1D**), as IHKA-L females spent more time in seizures than IHKA-L (p = 0.0007, **Figure 1D**) and IHKA-R males (p = 0.0004, **Figure 1D**), and IHKA-R females spent more time in seizures than IHKA-R males (p = 0.0007, **Figure 1D**).

**Figure 1.**
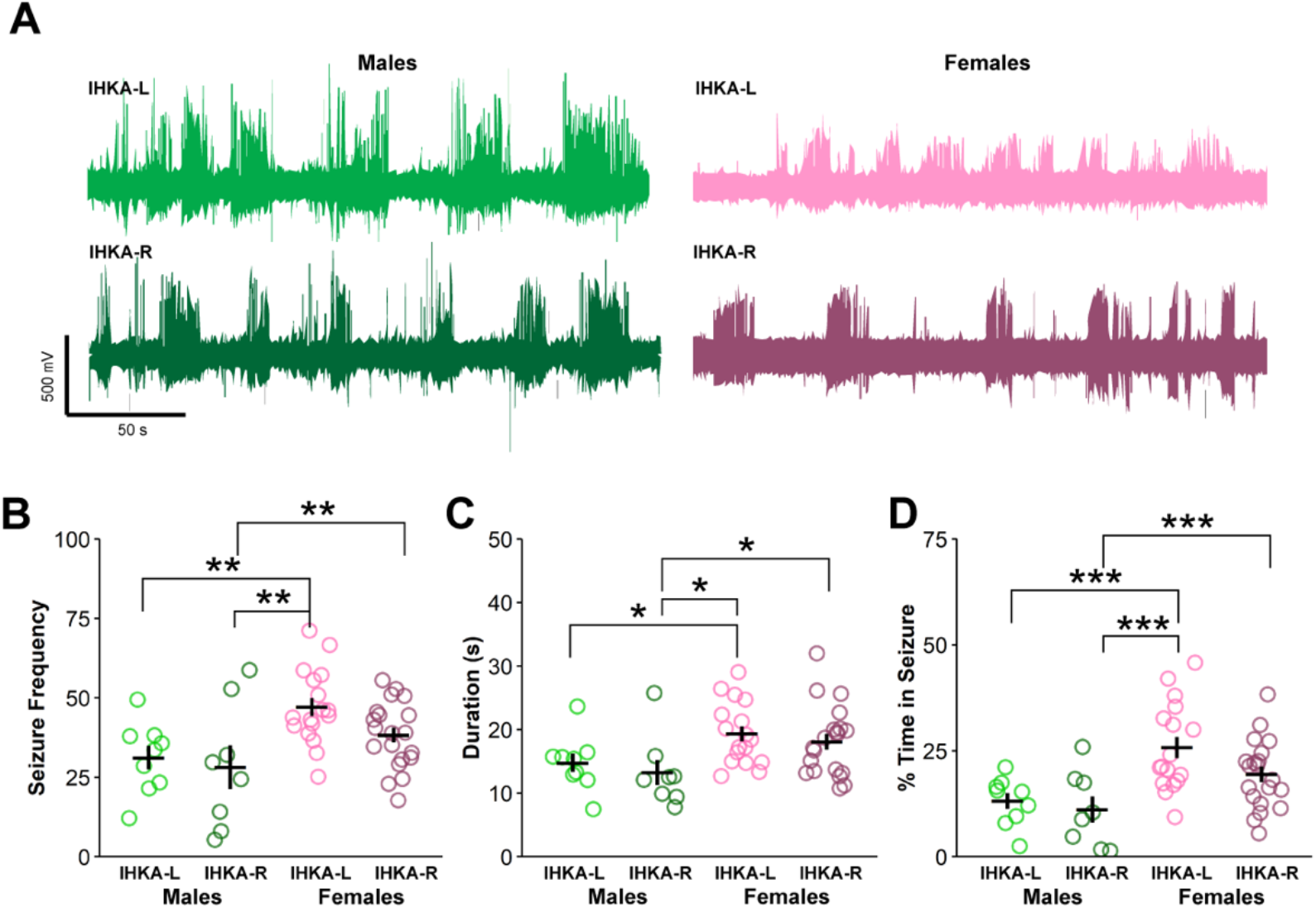
Sex differences in seizure parameters at 2 months after IHKA injection. A) Example seizure traces. B-D) Individual values of the average number of seizures per hour (B), average seizure duration (C), and average percent time spent in seizures (D) plotted with mean ± SEM. *p < 0.05, **p < 0.01, ***p < 0.001 two-way ANOVA and Tukey’s *post hoc* tests.

### Sex differences in seizure duration and percent time in seizures persist to 4 months after injection

To characterize the progression of seizure activity over time in these animals, the same animals evaluated at 2 mpi were also evaluated at 4 mpi, with the exception of two IHKA-R females that died prior to the 4-month time point. By 4 mpi, there was no effect of injection site (F(1,49) = 0.0001, p = 0.99, **Figure 2A**) or sex (F(1,48) = 2.17, p = 0.15, **Figure 2A**) on seizure frequency. Seizure duration, however, was still affected by sex and higher in females compared to males (F(1,49) = 11.84, p = 0.001, **Figure 2B**). Percent time spent in seizures was also influenced by sex (F(1,49) = 8.95, p = 0.004, **Figure 2B**), as IHKA females displayed higher percentage time in seizures than male counterparts (IHKA-L: p = 0.02, IHKA-R: p = 0.02, **Figure 2B**). Together, these results indicate that, compared to males, IHKA females maintain higher seizure duration and percent time spent in seizures as recorded at 4 mpi.

**Figure 2.**
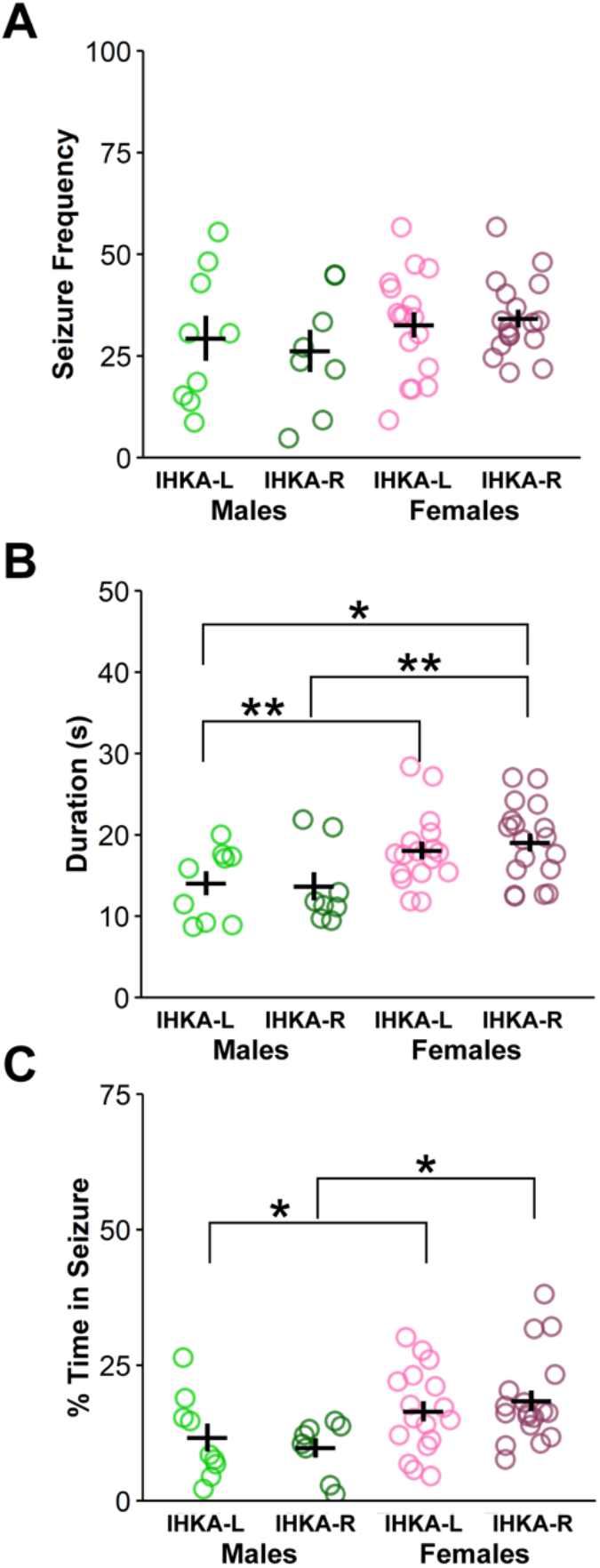
Higher seizure duration in females at 4 months after IHKA injection. A-C) Individual values of the average number of seizures per hour (A), average seizure duration (B), and average percent time spent in seizure (C) plotted with mean ± SEM. *p < 0.05, **p<0.01 two-way ANOVA and Tukey’s *post hoc* tests.

### Left-right asymmetry in seizure burden in IHKA females is most pronounced on diestrus at 2 months after injection

To investigate the importance of estrous cycle phase on seizure patterns, female animals were evaluated across diestrus, estrus, and proestrus. Recordings from three days of each estrous cycle stage from at least 3 different cycles per mouse were averaged to evaluate seizure parameters present on each estrous cycle stage. Evaluation of LFP recordings at 2 mpi showed that during diestrus and estrus, IHKA-L females displayed higher seizure frequency than IHKA-R (diestrus: t = 2.05, p-value = 0.05; estrus t = 2.41, p-value = 0.02, **Figure 3A**), with a trend towards elevated seizure frequency in IHKA-L females during proestrus (t = 1.97, p = 0.06, **Figure 3A**). However, seizure duration and percent time spent in seizures were higher in IHKA-L compared to IHKA-R females only during diestrus (seizure duration: W = 261, p-value = 0.04, **Figure 3B**; time spent in seizure: W = 286, p-value = 0.004, **Figure 3C**). These results suggest that there are distinct elevations in seizure parameters in IHKA-L compared to IHKA-R females at 2 mpi, and that these increases are more pronounced during diestrus.

**Figure 3.**
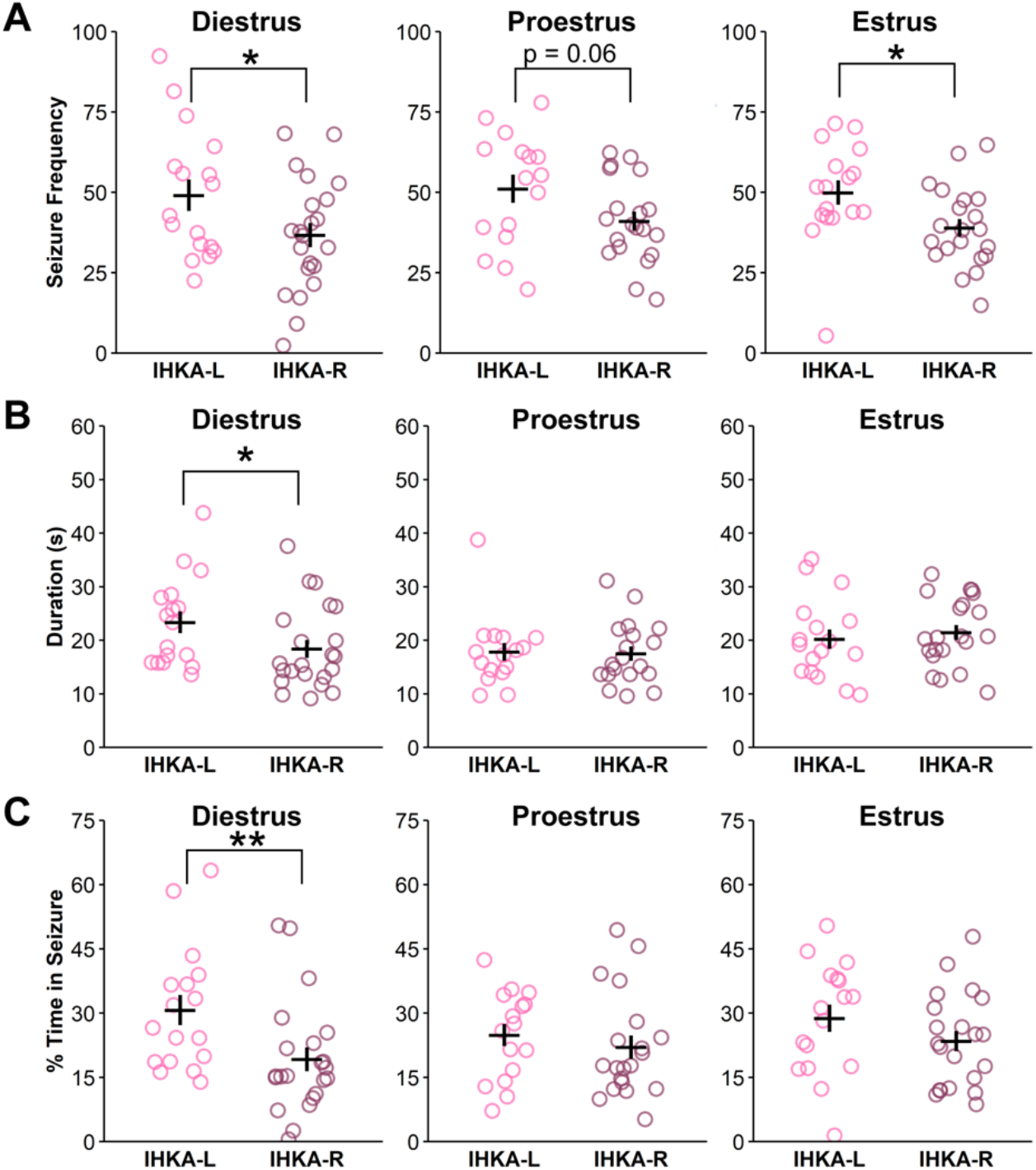
Seizure burden is elevated in IHKA-L females on diestrus 2 months after IHKA injection. A-C) Individual values of the average number of seizures per hour (A), average seizure duration (B), and average percent time spent in seizure (C) for IHKA females. Values are plotted with mean ± SEM. *p < 0.05, **p < 0.01 two-sample t-test or Wilcoxon ranked sum test based on data normality.

### Left-right asymmetries and effects of estrous cycle stage on seizure burden in females are dampened 4 months after injection

In contrast to the pattern of differences at 2 mpi, there were no appreciable differences between IHKA-L and IHKA-R females in any seizure parameters on diestrus or estrus at 4 mpi (**Figure 4**). During proestrus, however, IHKA-R animals showed trends for elevated seizure frequency (t = -2.00, df= 32, p = 0.05, **Figure 4A**) and percent time spent in seizures (t = -1.99, df = 32, p = 0.06, **Figure 4C**). These results suggest that there are less distinct influences of injection side on seizure parameters in females at 4 mpi.

**Figure 4.**
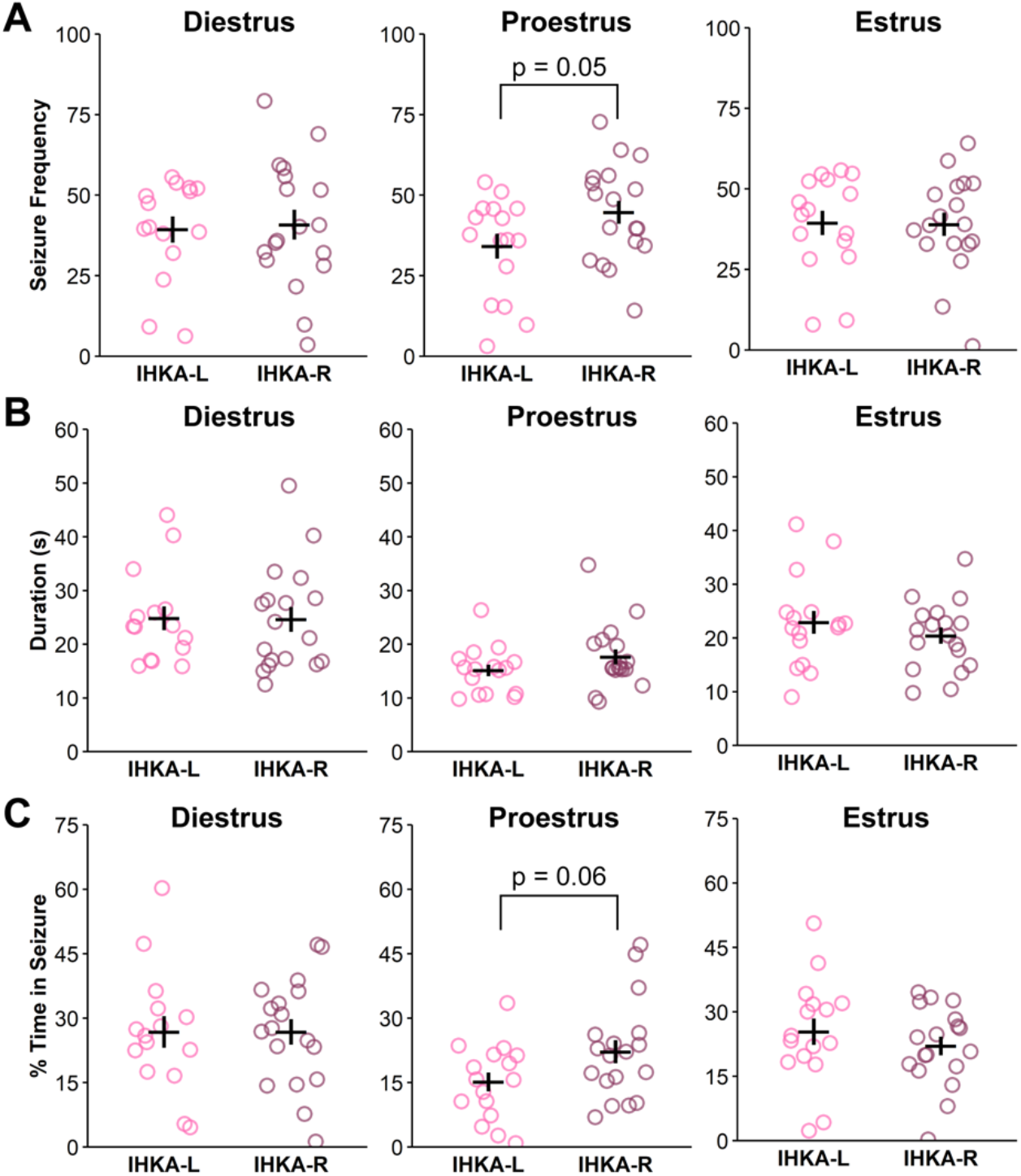
Lateralization and cycle stage effects on seizure burden in females are dampened at 4 months after IHKA injection. A-C) Individual values of the average number of seizures per hour (A), average seizure duration (B), and average percent time spent in seizure (C) for IHKA females. Values are plotted with mean ± SEM.

## Discussion

Clinical evidence indicates that the hemisphere of a temporal lobe seizure focus may influence seizure cluster patterning in women with epilepsy. Preclinical evidence suggests that the IHKA model of TLE holds validity for investigating mechanisms of the aforementioned clinical findings, as differences in hippocampal granule cell dispersion and pituitary gene expression are altered based on site of IHKA injection.^19^ Despite this evidence, the relationship between seizure patterns in IHKA animals and the site of injection has yet to be documented. In the present study we evaluated the development and presentation of seizures recorded with LFP at 2 and 4 mpi in male and female IHKA-L and -R mice. The results indicate a distinct sex difference in seizure parameters between male and female IHKA mice, with higher seizure burden in females. In addition, IHKA-L females display higher overall seizure burden on diestrus compared to IHKA-R females at 2 mpi. On proestrus and estrus, however, this effect is only present for seizure frequency. These results suggest that the phase of the estrous cycle influences left-right asymmetry in seizure burden in IHKA females.

Sex differences have been observed across multiple animal models of TLE.^27^ In the pilocarpine post-status epilepticus model of TLE, female rats typically have longer latency to acute seizures and lower mortality rates.^28^ In another study, male rats treated systemically with pilocarpine developed longer lasting seizures over a period of several months, whereas female rats tended to have higher frequency of seizures.^29^ In a systemic kainic acid injection model of TLE, male rats had greater susceptibility to seizures than their female counterparts.^28^ In mice, however, the results regarding systemic KA injection have been mixed, with one study reporting higher mortality rates, more severe seizures, and increased neurodegeneration in females,^30^ but another study describing higher mortality, seizure severity, cognitive impairment, hippocampal neuron loss, and reactive gliosis in males.^31^ With respect to the IHKA model, one report indicated that female mice do not show the same latent period duration following IHKA injection as males, and that hippocampal paroxysmal discharges were rare in females examined 4 weeks after injection.^32^ By contrast, another study conducted using the IHKA model did not indicate an effect of sex on seizures, cognitive impairment, or histopathology,^26^ although the latter study was not powered to evaluate sex differences. In the present study, males showed reduced overall seizure burden compared to females, which differs from other IHKA studies in mice. However, the effect of sex in animal models of epilepsy can vary greatly, and be influenced by factors such as methodology of seizure induction, measurement of seizures, the species of animal (or even strain), and the age at which seizures are induced.^27,33^

We evaluated seizure parameters on multiple stages of the estrous cycle to determine whether IHKA-L and IHKA-R mice show varied patterning across the different estrous cycle phases. On diestrus, proestrus, and estrus at 2 mpi, IHKA-L females showed higher seizure frequency in comparison to IHKA-R. With respect to seizure duration and percent time spent in seizures, however, IHKA-L mice only showed elevations on diestrus. Previous work established that the estrous cycle in rats can have impacts on interictal spike presentation in systemic kainic acid and pilocarpine models of TLE.^34^ Additionally, IHKA mice have previously been shown to have elevated seizure duration and time spent in seizures on proestrus and estrus combined compared to diestrus.^22^ The differences in seizure patterns of IHKA-L and -R mice on certain cycle stages may arise due to interactions between fluctuating ovarian hormones and basal differences in the left and right hippocampi. Such basal differences have been observed in mice as evidence has suggested that the left and right hippocampus have distinct populations of synapses,^35,36^ and that these differences contribute to left-right asymmetry in hippocampal function.^37^ There is also evidence that gonadal hormones can impact hippocampal lateralization, with androgen receptors suggested to underlie larger granule cell layer volume in the right hippocampus of both male and female C57BL/6J mice.^38^ Therefore, it is possible that the fluctuations in ovarian hormones across the estrous cycle, and lateralized interactions between ovarian hormones and the hippocampus, produce left-right asymmetry in emergent hippocampal phenotypes. In mice, the estradiol-to-progesterone (E:P) ratio is lower on diestrus than on proestrus and estrus.^27,39^ The variation in this hormone ratio across the mouse estrous cycle could thus contribute to the presentation of IHKA-L and IHKA-R seizure differences seen more prominently on diestrus compared to the other stages. However, previous studies have shown no difference in circulating estradiol and progesterone levels between IHKA-L and IHKA-R diestrous females.^19^

Human imaging studies suggest that brain networks can undergo reorganization across the menstrual cycle.^40,41^ In particular, fluctuations in progesterone that are common across the menstrual cycle have been associated with volumetric changes in hippocampal morphology.^42^ The impact of the menstrual cycle and ovarian hormones on the hippocampus may have functional ramifications as well, as worry and working memory can be affected by gonadal hormone levels.^43^ Specifically, higher levels of worry and lower working memory have been associated with increased estradiol and progesterone levels.^43^ The findings of the present study, in tandem with human imaging work, support the validity of mouse models in studies of mechanisms examining interactions between ovarian cycles and brain function and structure. Additionally, the presence of seizure burden asymmetries only in females aligns with clinical findings of reproductive endocrine disorder development among TLE patients. Male and female patients with TLE often develop reproductive endocrine dysfunction,^44,45^ but females more consistently develop higher rates of dysfunction than the general population.^45,46^ Moreover, the propensities for developing polycystic ovary syndrome or hypothalamic amenorrhea are differentially based on the hemisphere of the seizure focus in females.^47^ These findings indicate that there may be differential mechanisms in the left and right hemispheres that contribute to the ability of seizures to promote reproductive endocrine dysfunction. Furthermore, the different patterns of seizure clustering in association with menstrual cycle phase in females with left- or right-sided TLE^9,10^ suggest that the female brain may harbor left-right asymmetry that can shape the presentation of both epilepsy and associated comorbidities. The present findings of female-specific asymmetry are thus consistent with clinical reports, and support the translational validity of the IHKA mouse model in recapitulating at least some mechanistic differences in structural or functional changes due to seizures in the left and right hippocampus, as well as sex differences in this lateralization.

The variation in seizure patterning in mice when evaluated at 2 and 4 mpi indicates that seizure activity can shift with time after injection in females. Specifically, sex differences were pronounced by 2 mpi, but somewhat reduced by 4 mpi. In addition, IHKA-L females showed pronounced seizure burden elevations on diestrus at 2 mpi that appeared to abate by 4 mpi. Trends toward elevated seizure frequency burden were observed in IHKA-R mice on proestrus at 4 mpi, in contrast to the data obtained at 2 mpi. Our group has shown that most IHKA female mice exhibit elongated estrous cycles.^19,22–24,48,49^ In our previous work, estrous cycle disruption persisted to 4 mpi in IHKA-R but not IHKA-L females.^19^ The dissipation in phenotypes in IHKA-L females suggests that asymmetries may be influenced by the progression of the disease, the increased age of the animals, or both. The progressive decline of certain physiological functions as mice age could shape the varied patterns seen at different timepoints. In total, these findings suggest that injection site and time of recording after injection in IHKA female mice are two factors that should be considered in experimental design and can shape overall seizure burden outcomes.

The IHKA mouse model displays many characteristics recapitulating human epilepsy, underscoring the validity of its use in investigating neural mechanisms of TLE. The present work indicates that seizure burden outcomes within this preclinical model are influenced by sex and, in females, laterality of epileptogenic insult. Female mice also show estrous cycle-associated changes in seizure activity that vary depending on the hemisphere of targeted hippocampal damage. This pattern likely reflects hippocampal lateralization, compounded by the influence of physiological fluctuations across the estrous cycle. This work thus reveals female-specific left-right asymmetry in functional properties of the murine hippocampus, with implications for both normal and disease states.

## Acknowledgements

We thank Jiang Li for assistance with pilot studies and Victoria Daniels for help with estrous cycle monitoring. This work was supported by the National Institutes of Health (NIH)/National Institute of Neurological Disorders and Stroke and the NIH Office of Research on Women’s Health through grants R03 NS103029 and R01 NS105825 (C.A.C.-H.) and by a CURE Epilepsy Research Continuity Fund Grant (C.A.C.-H.).

## Author contributions

C.A.C.-H. contributed to the conception and design of the study; C.A.C. and L.K.L. contributed to the acquisition of data; C.A.C. contributed to the analysis of data; C.A.C. and C.A.C.-H. contributed to drafting the manuscript and preparing the figures

